# Genotype and dose-frequency may critically determine the therapeutic efficacy of chronic oxytocin treatment in humans

**DOI:** 10.1101/493387

**Authors:** Juan Kou, Yingying Zhang, Feng Zhou, Cornelia Sindermann, Christian Montag, Benjamin Becker, Keith M Kendrick

**Affiliations:** The Clinical Hospital of Chengdu Brain Science Institute, MOE Key Laboratory for Neuroinformation, University of Electronic Science and Technology of China, Chengdu 611731, China; Department of Molecular Psychology, Institute of Psychology and Education, Ulm University, Ulm, Germany

## Abstract

Chronic intranasal oxytocin administration daily is increasingly proposed as a therapy for social dysfunction but some clinical trials have reported small or no beneficial outcomes. No empirical evidence proves that this is optimal therapeutically or whether oxytocin receptor genotype influences treatment sensitivity. In a randomized, placebo-controlled pre-registered trial on 138 adult male subjects we investigated effects of single and repeated oxytocin treatment (24IU daily or alternatively days for 5 days). Primary neural outcomes assessed core therapeutic mechanisms of action, i.e. amygdala fear reactivity and amygdala-prefrontal intrinsic functional connectivity and modulation by oxytocin receptor polymorphisms (rs53576, rs2254298). The expected oxytocin-induced reduction in amygdala fear reactivity and associated anxious-arousal following single-dose administration was abolished after daily treatment but maintained when administered every other day. Oxytocin selectively reduced amygdala and arousal fear reactivity in AA homozygotes of rs53576 and A+ carriers of rs2254298. By contrast, oxytocin-enhanced intrinsic amygdala-prefrontal coupling was maintained independent of dose frequency and genotype. Together the findings provide the first evidence that infrequent rather than daily oxytocin administration protocols may be therapeutically most efficient and that its neural and behavioral anxiolytic actions are highly genotype-dependent.

## Introduction

Intranasal oxytocin (OT) has been proposed as novel treatment to attenuate social dysfunctions and anxiety in autism spectrum disorder, social anxiety and schizophrenia (Kendrick et al., 2017; Meyer-Lindenberg et al., 2011; Young and Barrett, 2015) as well as an augmentative strategy to facilitate fear extinction (Eckstein et al., 2015) and working memory performance (Zhao et al., 2018). Preclinical studies consistently suggest that effects on amygdala functioning (attenuated fear reactivity, increased amygdala-prefrontal intrinsic connectivity, (Eckstein et al., 2015; Meyer-Lindenberg et al., 2011) represent the primary therapeutic-relevant neural mechanism of action of intranasal OT. The proposed neural mechanisms have been validated using single dose administration protocols (with 24 International Units (IU) being established as optimal (Spengler et al., 2017)), however, in contrast initial clinical trials with chronic (twice daily) administration protocols over longer intervals reported inconsistent, or at best modest therapeutic efficacy on social-emotional dysfunctions (Guastella et al., 2015; Halverson et al., 2019; Jarskog et al., 2017; Kendrick et al., 2017; Watanabe et al., 2015; Yamasue et al., 2018).

G-protein-coupled neuropeptide receptors typically exhibit internalization following even a relatively short period of constant exposure to their target peptides and cannot subsequently respond (desensitized) until they are recycled back the surface of the cell membrane (Pierce et al., 2002; Lohse and Hofmann, 2015). Indeed, neuroendocrine neurons in the hypothalamus regulating pituitary peptide release tend to exhibit phasic discharge patterns resulting in a pulsatile pattern of release which may serve to reduce internalization of their target receptors (Russell, 2018). Growing evidence from both in vitro and animal studies suggest there may be extensive internalization and subsequent desensitization of OT receptors (OXTR) following chronic OT administration (Smith et al., 2006; Stoop, 2012). In terms of neural OXTR receptors those in the amygdala may be particularly susceptible to desensitization (Terenzi and Ingram, 2005) and this region critically mediates many of the effects of intranasal OT on social cognition and anxiety (Kendrick et al., 2017). Moreover, rodent models have reported that in contrast to single doses, chronic administration of OT can actually produce social impairment (Bales et al., 2013; Du et al., 2017; Huang et al., 2014) in the context of reduced receptor expression in the amygdala and nucleus accumbens (Du et al., 2017). Furthermore, an initial study reported that chronic versus single administration of OT results in divergent neurochemical changes in both the rodent and human frontal cortex (Benner et al., 2018), emphasizing high translational relevance of the preclinical animal models for clinical trials employing chronic treatment protocols. Thus, empirically evaluated optimal treatment protocols for chronic intranasal OT administration in humans are urgently needed to determine the therapeutic potential of OT for social-emotional dysfunctions in psychiatric disorders.

A further unresolved issue which may impede the therapeutic efficacy of intranasal OT is that different OT receptor (OXTR) genotypes have been associated with social behaviors and may also influence sensitivity of behavioral and neural responses to intranasal OT. In particular, OXTR polymorphisms rs53576 and rs2254298 have been associated with autism (Cataldo et al., 2018) and deficits in social and emotional processing and anxiety (Jurek and Neumann, 2018; Parker et al., 2014; Yang et al., 2017) as well as individual variations in behavioral and neural responses to intranasal OT (Chen et al., 2015; Feng et al., 2015a). Indeed, variants of rs2254298 may represent a trans-diagnostic biomarker for social dysfunctions (Brüne, 2012). It is therefore imperative to establish whether OXTR polymorphisms critically determine sensitivity to the key neural mechanisms of intranasal OT in order to identify treatment-responsive individuals likely to exhibit therapeutic benefits.

Although single doses of OT have been reported to produce numerous functional effects, those on amygdala functioning have been most consistently determined as promising key neural mechanisms: attenuated amygdala fear responses and increased resting state functional connectivity between the amygdala and medial prefrontal cortex (Kendrick et al., 2017; Spengler et al., 2017). Both neural markers have been primarily associated with attenuated anxiety in terms of reduced responses to social threat and enhanced top-down control of emotion (Zhao et al., 2019), although OT effects on the amygdala may also mediate its actions on a range of social cognition domains (Kendrick et al., 2017; Meyer-Lindenberg et al., 2011; Spengler et al., 2017; Young and Barrett, 2015).

In order to determine optimal treatment protocols for chronic intranasal OT administration the current pre-registered double-blind, randomized between subject placebo (PLC)-controlled pharmacological neuroimaging trial (see Fig. 1) in 138 adult male subjects therefore aimed at determining the effects of acute (single dose) and repeated doses (daily or every other day for 5 days) of 24IU OT versus PLC on amygdala-centered neural and behavioral (valence, arousal and intensity ratings of fear faces) mechanisms of OT. To identify individuals with the most promising treatment sensitivity we additionally assessed the influence of OXTR rs53576 and rs2254298 polymorphisms on acute and repeated dose OT effects.

**Fig.1.**
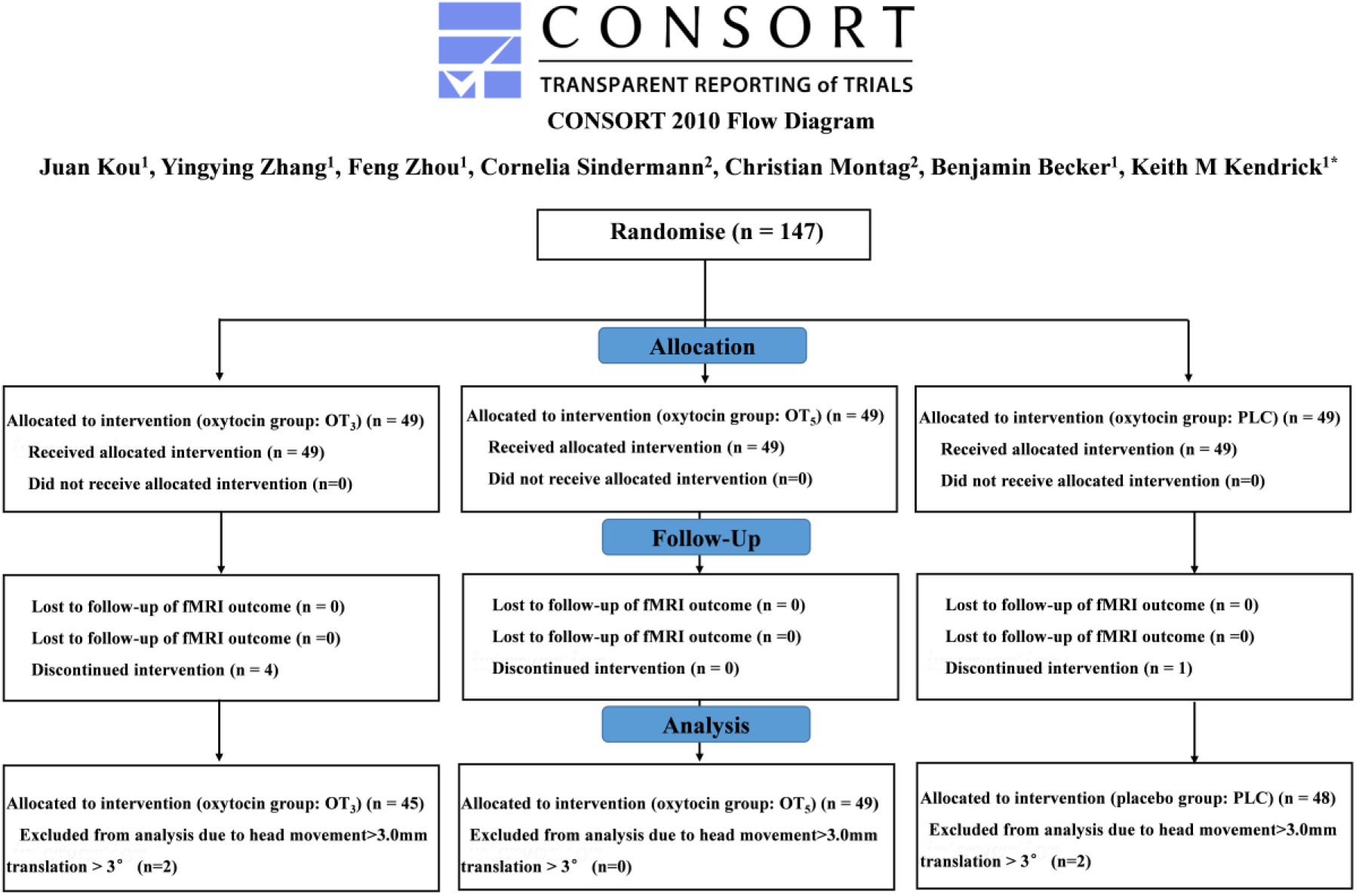
CONSORT flow diagram of the clinical trial.

## Results

### Acute effects of OT on neural responses to emotional faces

In accordance with the primary outcome measure of the trial analyses focused on the neural responses to fearful faces, however no significant effects of OT were found for happy or angry faces. Comparison of single-dose OT-(combined OT_3_ and OT_5_ groups) and PLC-treated subjects (1^st^ day) revealed significantly decreased right amygdala reactivity towards fearful faces on the whole brain level (k = 78, p_FWE =_ 0.043, x=27, y=-4, z=-13) (Fig.2A). Cytoarchitectonic probabilistic localization (Anatomy toolbox V1.8 (Eickhoff et al., 2005)) mapped the peak coordinate with >80% probability to the basolateral amygdala sub-region. In addition, OT also decreased fear-reactivity in the right superior frontal gyrus and bilateral primary visual cortex (see **SI** and Table S1). There were no significant differences between the OT_3_ and OT_5_ groups on the 1^st^ day.

**Fig.2.**
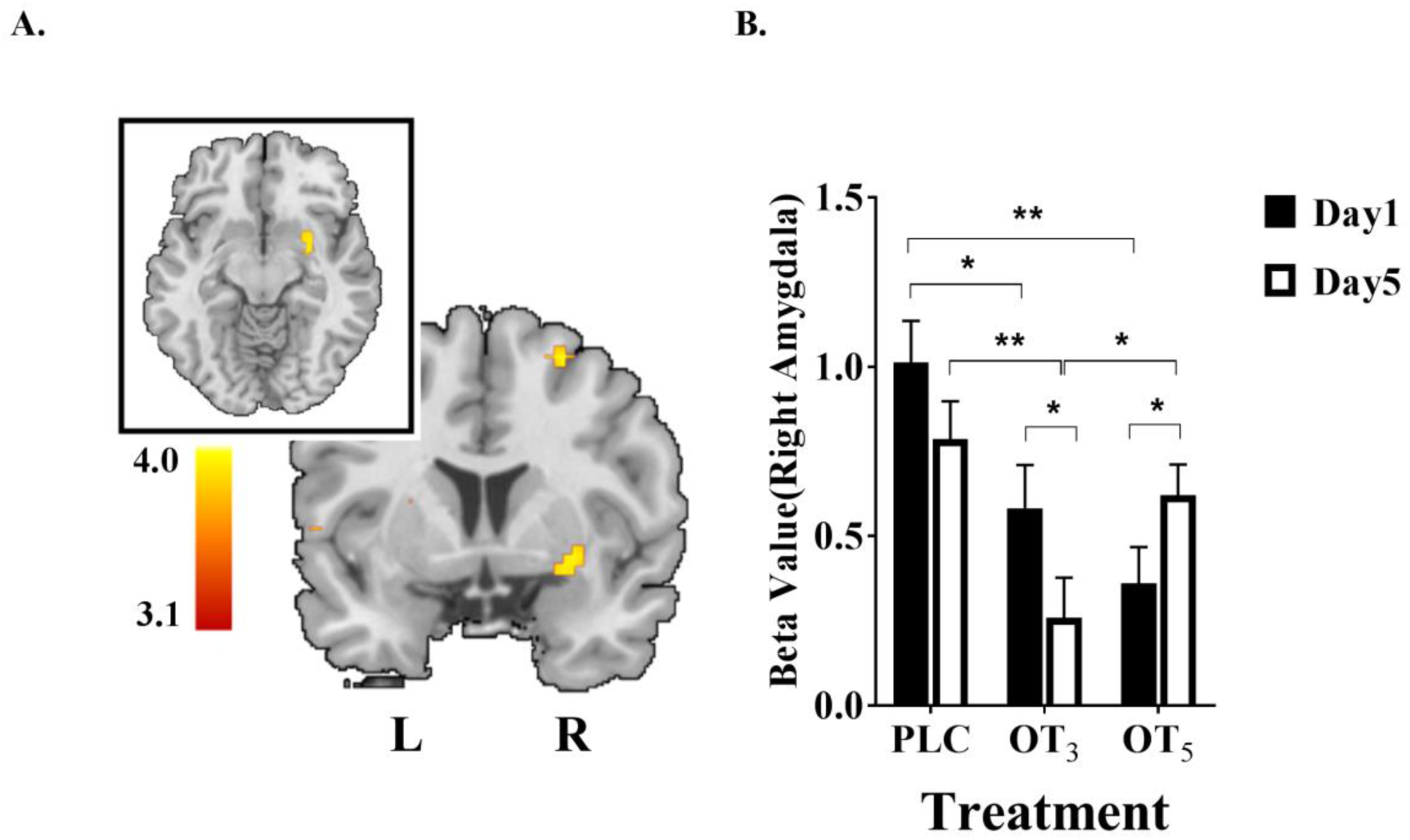
The effect of intranasal oxytocin (OT) treatment on neural responses to fearful faces on the 1st day. (A) The t-map of the treatment effect (FWE c = 78, t = 3.93, p = 0.043) showed an activated cluster peaking at the right amygdala (x = 27, y = −4, z = −13). (B) Parameter estimates extracted using a 6-mm radius sphere centered at the peak MNI coordinates at the right amygdala on the 1^st^ and 5^th^ day separately revealed that OT treatment on alternate days (OT_3_) decreased amygdala responses on both the 1^st^ and 5^th^ days whereas for the daily OT treatment group (OT_5_) group the decrease only occurred on the 1^st^ day. * p < 0.05, ** p < 0.01, two-tailed t-test. Bars indicate M ± SE.

### Primary outcome measures: effects of repeated doses on OT-evoked changes

Mixed ANOVAs with treatment (PLC, OT_3,_ OT_5_) and time point (1^st^ day, acute effects; 5^th^ day, chronic effects) as factors and right amygdala fear reactivity as dependent variable revealed a significant main effect of treatment (F_2, 135_ = 7.85, p = 0.001, *η*^2^_p_ = 0.104) and a treatment x time point interaction (F_2, 135_ = 5.74, p = 0.004, *η*^2^_p_ =0.078). Post-hoc Bonferroni corrected comparisons between groups on the 5^th^ day revealed that the OT_3_ group exhibited a suppression of right amygdala fear reactivity relative to both the PLC (p = 0.001, Cohen’s *d* = 0.70, 95% CI, −0.833 to −0.224) and OT_5_ groups (p = 0.018, *d* = 0.52, 95% CI, −0.662 to −0.062) but the OT_5_ group did not (p = 0.266, relative to the PLC group). Within group comparisons showed that whereas amygdala reactivity did not change on day 1 relative to day 5 in the PLC group (p = 0.090, *d* = 0.29, 95% CI, −0.035 to 0.487), its suppression was significantly enhanced in the OT_3_ group (p = 0.019, *d* = 0.40, 95% CI, 0.054 to 0.595) and attenuated in the OT_5_ one (p = 0.044, *d* = 0.38, 95% CI, −0.514 to −0.007) (Fig. 2B).

For the behavioral ratings mixed ANOVAs with treatment and time point as within-subject factors revealed a significant main effect of treatment for arousal, but not intensity ratings (arousal: F_1, 135_ = 4.99, p = 0.008, *η*^2^_p_ = 0.07, intensity: F_1, 135_ = 2.60, p = 0.078, *η*^2^_p_ = 0.04), and time point for both emotional arousal and intensity ratings (arousal: F_1, 135_ = 12.09, p = 0.001, *η*^2^_p_ = 0.082, intensity: F_1, 135_ = 10.81, p = 0.001, *η*^2^_p_ = 0.74) although no significant two-way interactions (arousal: F_1, 135_ = 1.85, p = 0.162, *η*^2^_p_ = 0.03, intensity: F_1, 135_ = 2.97, p = 0.055, *η*^2^_p_ = 0.04). An exploratory post hoc analysis showed that both emotional arousal and intensity ratings for fear faces in the OT_3_ group were decreased on the 5^th^ day (arousal: p = 0.001 versus PLC, *d* = 0.63, 95% CI, −1.544 to −0.420, p = 0.033 versus OT_5_, *d* = 0.42, 95% CI, −1.158 to −0.051; intensity: p = 0.004 versus PLC, 95% CI, −1.299 to −0.247, *d* = 0.87, p = 0.088 versus OT_5_, *d* = 0.35, 95% CI, −0.968 to 0.068). Although there was a significant decrease in arousal ratings in the OT_3_ group on the 1^st^ day compared to the PLC group there was no difference between the OT_3_ and OT_5_ groups (arousal: p = 0.015 versus PLC, *d* = 0.31, 95% CI, 0.129 to 1.175, p = 0.355 versus OT_5_, 95% CI, −0.268 to 0.744). If we combined the two OT groups on the 1^st^ day to increase statistical power there was a marginal effect of decreased arousal but not intensity ratings (arousal: p = 0.060 versus PLC, *d*=0.35; intensity p = 0.150) (Fig.3). No significant main effects or interactions were found for valence ratings (all ps > 0.120).

**Fig.3.**
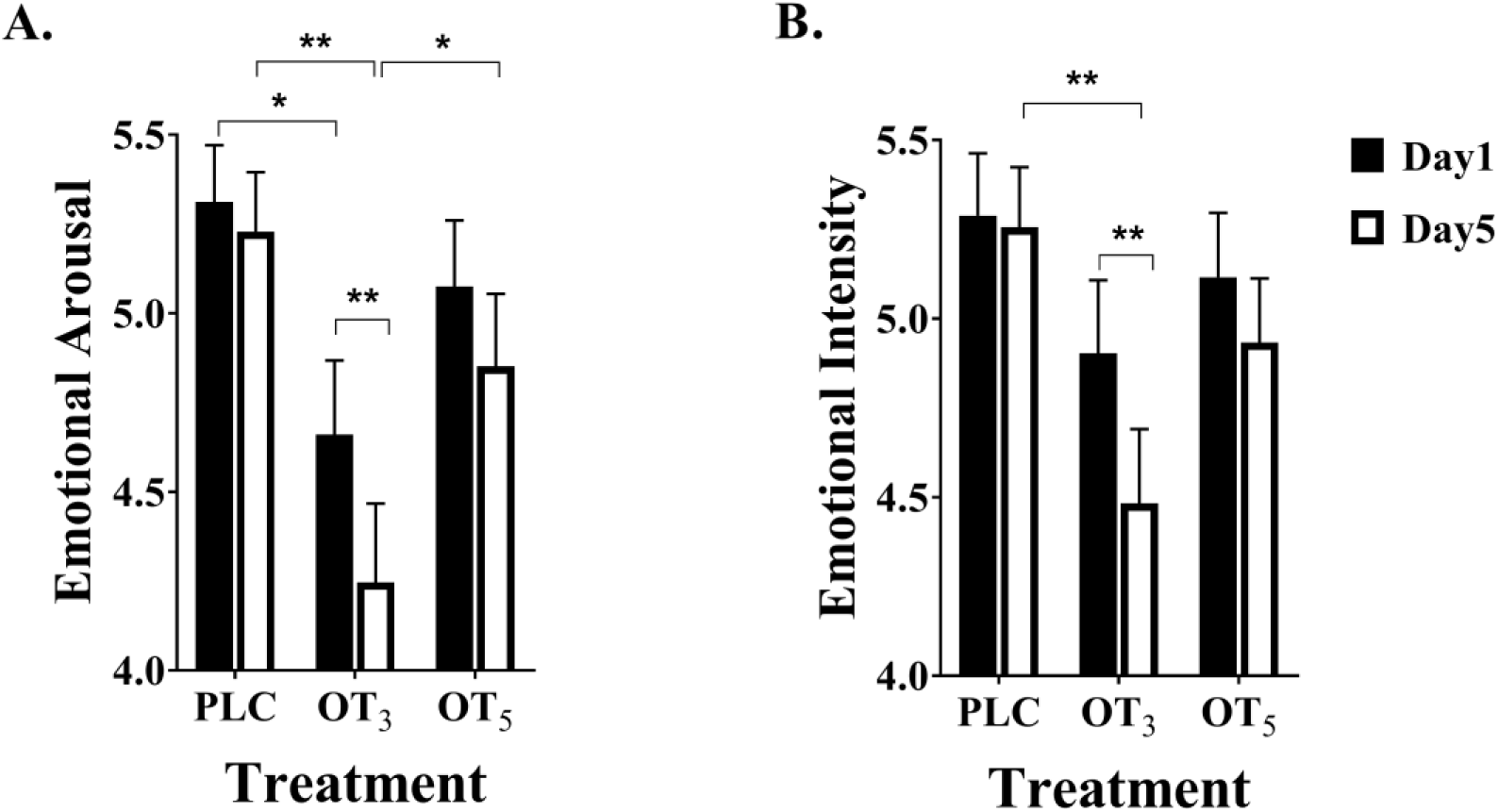
Oxytocin influenced intensity and arousal ratings. Oxytocin decreased intensity and arousal ratings of fearful faces only on the 5^th^ day of treatment in the group receiving OT on alternate days (OT_3_) compared with placebo (PLC). *p< 0.05. **p<0.01. Bars depict M ± SE.

Although not included in our primary outcome measures the (calcarine) visual cortex showed the same reduced responsivity to daily OT as the amygdala, and a similar pattern of association with behavioral intensity and arousal ratings, whereas the superior frontal gyrus did not (see **SI**). Voxel based morphometry analysis revealed no evidence for acute or repeated dose effects of OT on gray matter volume (see **SI**).

### Associations between amygdala responses and behavioral ratings to fear faces

Significant associations between right amygdala activation and emotional arousal and intensity scores were observed in the PLC group on the 1^st^ day (arousal: r = 0.44, p = 0.003; intensity: r = 0.54, p < 0.001) demonstrating that greater amygdala activation by fear faces was associated with increased anxiety. This association was absent in both OT groups and significantly different from the PLC group on the 1^st^ day (arousal: OT_3_, r = −0.21 p = 0.181; Fisher’s z = 3.08, p = 0.002; OT_5_, r = 0.09, p = 0.555; z = 1.79, p = 0.073; intensity: OT_3_, r = −0.13 p = 0.399; Fisher’s z = 3.33, p < 0.001; OT_5_, r = 0.12, p = 0.413; z = 2.35, p = 0.024). The same effect was also seen on the 5^th^ day for arousal ratings (PLC: r = 0.33 p = 0.024; OT_3_, r = −0.21 p = 0.187; z = 2.52, p = 0.012; OT_5_ group, r = −0.15 p = 0.317; z = 2.33, p = 0.020) although slightly weaker for intensity ratings (PLC: r = 0.25 p = 0.092; OT_3_, r = −0.29 p = 0.057; z = 2.53, p = 0.011; OT_5_ group, r = −0.03 p = 0.847; z = 1.34, p = 0.180) (Fig.4).

**Fig.4.**
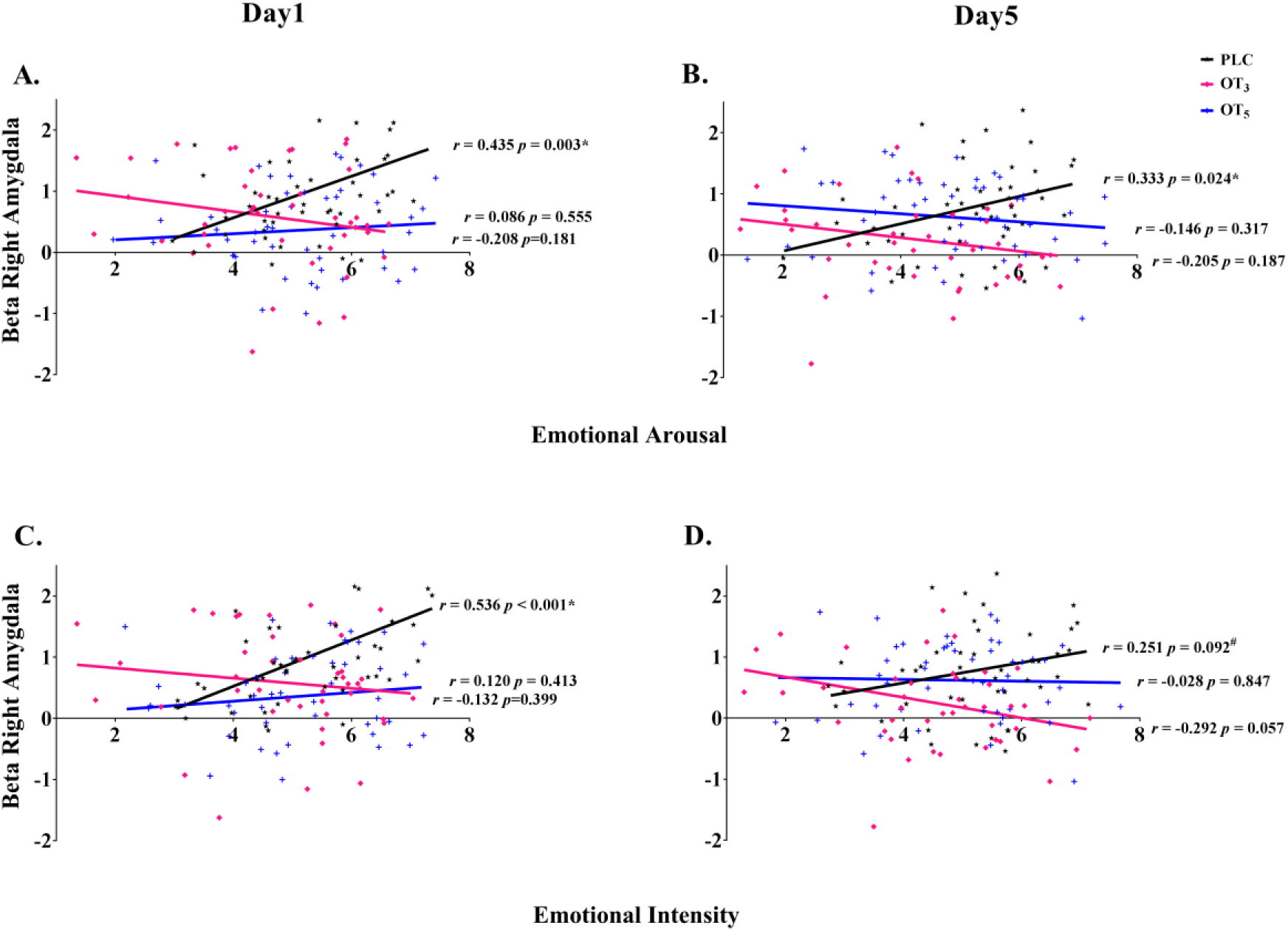
Associations between intensity and arousal ratings of fear faces and amygdala activation in the three treatment groups (PLC, OT_3_ and OT_5_) on days 1 and 5. *p< 0.05. **p<0.01

### Effects of repeated oxytocin doses on resting-state functional connectivity

Right-amygdala seed-to-whole brain fMRI resting-state analysis by mixed-effect ANOVA revealed a main effect of treatment (OT_3_, OT_5_ and PLC) on amygdala functional coupling with vmPFC (peak MNI x = −3, y =53, z = −19, F_2, 135_ = 16.55, p_FWE_ = 0.001, k = 88) before subjects underwent the face paradigm, whereas no brain regions showed a significant time point x treatment interaction. Post-hoc analyses demonstrated that right amygdala intrinsic connectivity with the vmPFC was stronger in both the OT_3_ (k = 74, p_FWE_ = 0.014, x = −3, y = 53, z = −19) and OT_5_ groups (k = 127, p_FWE_ = 0.001, x = −3, y = 53, z = −19) relative to PLC. Subsequent confirmatory analyses of resting state functional connectivity in this pathway acquired after the face task paradigm also revealed a significant main effect of treatment (F_2, 135_ = 7.08, p = 0.001, *η*^2^_p_ = 0.095). Post hoc comparisons showed both groups exhibited increased connectivity in this pathway after a single OT dose on the 1^st^ day (OT_3_, p = 0.001, *d* = 0.71; OT_5_, p = 0.063, *d* = 0.40; relative to PLC) and after repeated doses on the 5^th^ day (OT_3_ p = 0.047, *d* = 0.46; OT_5_, p = 0.001, *d* = 0.69; relative to PLC). The OT_3_ and OT_5_ groups did not differ significantly on the 1^st^ (p = 0.140) or 5^th^ (p = 0.208) day (Fig. 5B).

**Fig.5.**
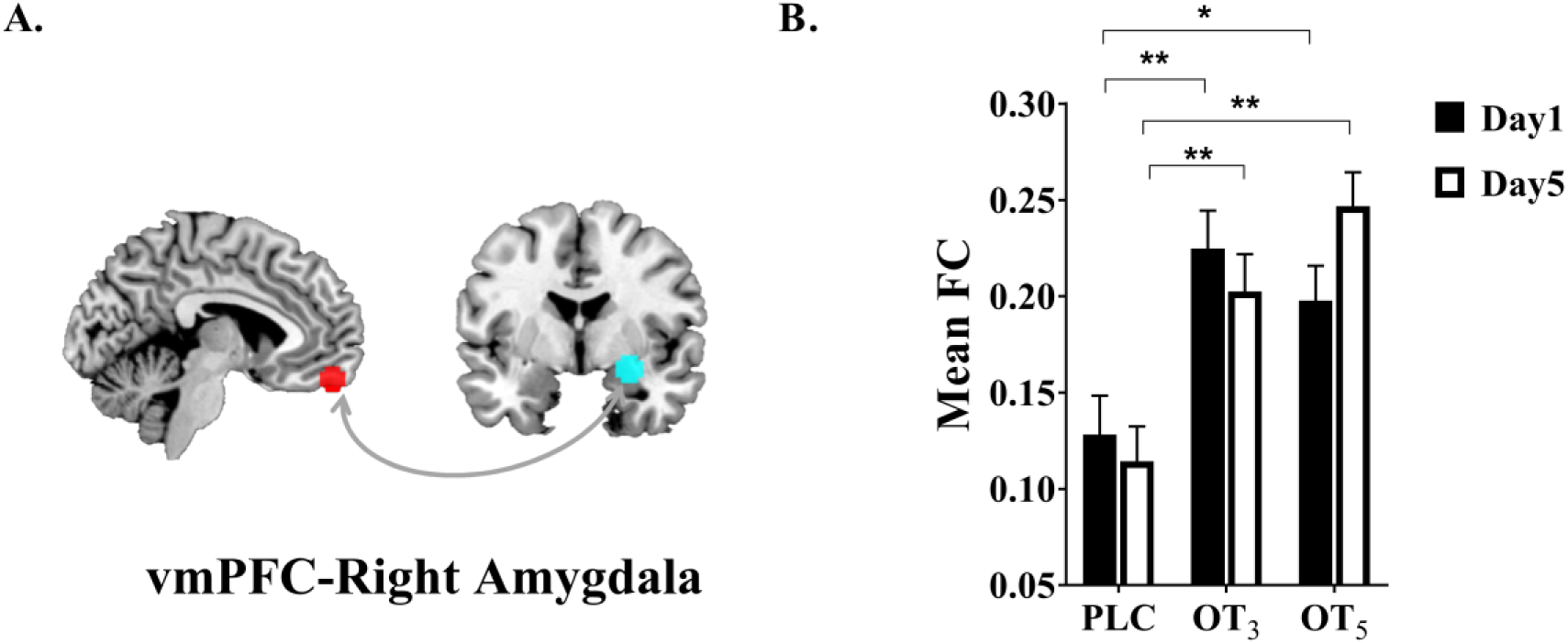
The effects of oxytocin (OT) on resting state functional connectivity (A) schematic showing functional connection between the right amygdala and ventromedial prefrontal cortex (vmPFC). Whole brain functional connectivity (FC) used the right amygdala as a region of interest (6mm sphere, x=27, y=-4, z=-13) and (B) revealed significant increased functional connectivity between vmPFC and right amygdala (p_FWE_ <0.05, x=-3, y=53, z=-19) in both OT treatment groups (every day – OT and every other day - OT on both days 1 and 5). Data for the two resting state periods before and after the face emotion task were similar and are therefore showed here combined. Bar graph illustrates the extraction of parameter estimates from right amygdala connectivity with vmPFC (M ± SE). * p < 0.05, ** p < 0.01, two-tailed t-test. Bars indicate M ± SE.

### Associations with OXTR genotype

The number of G-carriers and G-non-carriers of rs53576 as well as A-carriers and A-non-carriers of rs2254298 did not differ between the three groups and both SNPs satisfied the Hardy Weinberg Equilibrium (see Tables S2 **and** S3). With Bonferroni-correction for the multiple SNPs and alleles (i.e. 2 x 2 = 4), p < 0.0125 was considered significant. For amygdala responses there was a significant treatment x genotype interaction for rs53576 (F_2, 105_ = 6.01, p = 0.003, *η*^2^_p_ = 0.10). Post-hoc analysis revealed that OT-induced amygdala suppression on the 1^st^ day was only significant in G-non-carriers (p < 0.001, *d* = 0.99, 95% CI, 0.432 to 1.333) and also on the 5^th^ day in the OT_3_ group (p < 0.001, *d* =1.41, 95% CI, 0.638 to 1.483). While the interaction between treatment and genotype did not achieve significance for rs2254298 (F_2, 105_= 1.48, p = 0.232, *η*^2^_p_ = 0.03) an exploratory post hoc analysis revealed that OT-induced suppression on the 1^st^ day across treatment groups (OT_3_ and OT_5_ groups combined to increase power) was only significant in A-carriers (p = 0.001, *d* = 0.91, 95% CI, 0.337 to 1.244) and also on the 5^th^ day in the OT_3_ group (p = 0.004, *d* =0.81, 95% CI, 0.218 to 1.122) (see Fig. 6). Similar patterns of genotype association were found for the visual cortex but not superior frontal gyrus responses to fear faces (see **SI**).

**Fig.6.**
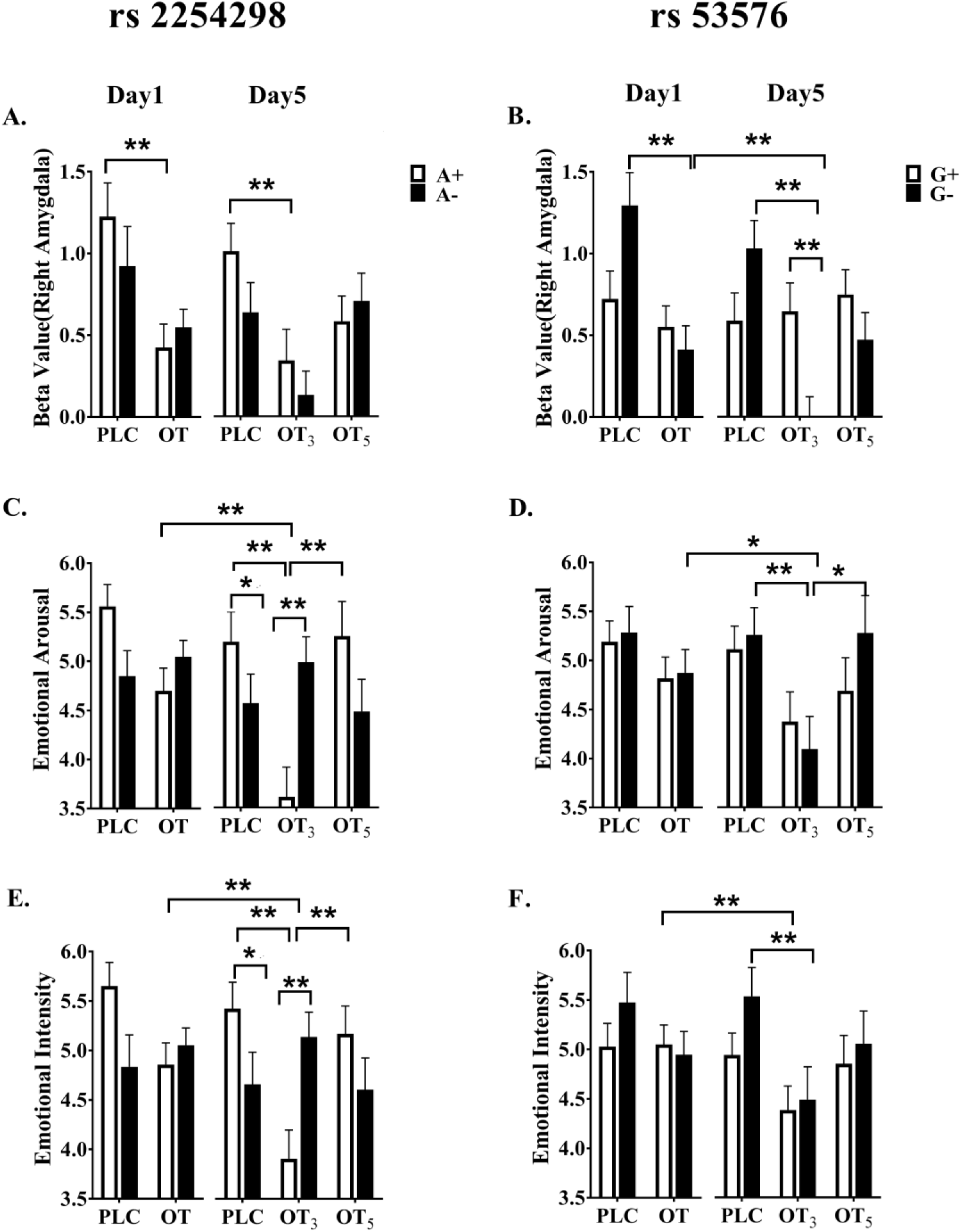
Influence of oxytocin receptor genotype on right amygdala responses to fearful faces (n = 111 subjects). (A)-(B): for rs 53576 only G-non carriers (i.e. AA) showed reduced amygdala response to fearful faces and for rs2254298 only A-carriers (i.e. AA and AG) in the groups with intranasal oxytocin treatment either daily (OT_5_) or every other day (OT_3_). (C)-(D): A similar pattern was found for rs2254298, but not rs53576 for arousal ratings of fearful faces although only on day 5 in the OT_3_ group. (E)-(F): A similar pattern was found for rs2254298, but not rs53576 for intensity ratings of fearful faces although only on day 5 in the OT_3_ group ** p < 0.005, * p < 0.0125, two-tailed t-test (Bonferroni corrected significance threshold of p = 0.0125). Bars indicate M ± SE.

For both arousal and intensity ratings there was a significant treatment x genotype interaction for rs2254298 (arousal: F_2, 105_= 9.70, p < 0.001, *η*^2^_p_ = 0.16, intensity: F_2, 105_= 7.42, p = 0.001, *η*^2^_p_ = 0.12). To increase the statistical power, we combined the OT_3_ and OT_5_ groups on the 1^st^ day. Post hoc analysis showed a significant decrease in ratings on the 1^st^ day in the OT group only in A-carriers (arousal: p = 0.014 versus PLC, *d* = 0.64, 95% CI, 0.177 to 1.542; intensity: p = 0.024 versus PLC, *d* = 0.60, 95% CI, 0.109 to 1.478). There was a significant decrease in ratings on the 5^th^ day in the OT_3_ group only in A-carriers (arousal: p < 0.001 versus PLC, *d* = 1.69, 95% CI, −2.780 to −1.231; p < 0.001 versus OT_5_, *d* = 1.10, 95% CI, −2.414 to −0.865; intensity: p < 0.001 versus PLC, *d* = 1.56, 95% CI, −2.531 to −1.075; p = 0.001 versus OT_5_, *d* = 0.97, 95% CI, −1.993 to −0.536). There was also a significant difference between the 1^st^ and 5^th^ days in the OT_3_ group only in A-carriers (arousal: p=0.002, *d* =1.72, 95% CI, 0.201 to 0.862, intensity: p<0.001, *d*=0.42, 95% CI, 0.272 to 0.914) not the other two groups (ps > 0.06) and there was a significant difference between A-carriers and A-non carriers in the OT_3_ group (arousal: p=0.001, *d* =1.07, 95% CI, 0.578 to 2.172, intensity: p=0.001, *d*=1.01, 95% CI, 0.483 to 1.982) and in PLC group (arousal: p=0.032, *d* =0.92, 95% CI, −1.783 to −0.084, intensity: p=0.021, *d* = 0.88, 95% CI, −1.742 to −0.144) on the 5^th^ day but not in OT_5_ (ps>0.088)

There was no significant treatment x genotype interaction for rs53576 (arousal: F_2, 105_= 1.28, p = 0.283, *η*^2^_p_ = 0.02, intensity: F_2, 105_= 0.52, p = 0.599, *η*^2^_p_ = 0.01). However, an exploratory post hoc analysis found that the reductions in arousal and intensity ratings in the OT_3_ group were only significant in G-non carriers on the 5^th^ day (arousal: p = 0.006 versus PLC, *d* = 0.87, 95% CI, −1.980 to −0.348; p = 0.012 versus OT_5_, *d* = 0.84, 95% CI, −2.102 to 0.265; intensity: p = 0.008 versus PLC, *d* = 0.72, 95% CI, −1.810 to −0.285; p = 0.193 versus OT_5_) (Fig. 6).

Since comparable patterns of OT effects were observed on pre- and post-task resting state data right amygdala-vmPFC functional connectivity values were pooled to increase power (Fig. 4B). However, no significant main or interaction effects of genotype for either SNP (rs 2254298 all ps > 0.262; rs53576 all ps > 0.396) were found.

## Discussion

Overall, our findings firstly validate the use of OT-reductions in amygdala responses to fear-faces and associated emotional intensity and arousal ratings together with increased resting state functional connectivity between the amygdala and vmPFC as robust markers for its putative therapeutic mechanisms of action. Intriguingly, these task-dependent and resting-state effects of OT show a markedly different sensitivity to repeated intranasal doses and OXTR genotype, although our results indicate that to achieve both maximal task and resting-state changes an optimal protocol using the standard 24IU dose may be to administer it every other day rather than daily. Indeed, both amygdala and behavioral anxiolytic responses to fearful faces were even more pronounced after 5 days when OT was administered every other day and with resting-state functional connectivity changes there was no advantage in terms of their magnitude when OT was administered daily as opposed to every other day.

Our findings are therefore highly consistent with preclinical animal models demonstrating OXTR desensitization following repeated doses of OT in some brain regions (Terenzi and Ingram, 2005; Smith et al., 2005; Stoop, 2012) and that chronic administration can reduce brain OXTR expression (Du et al., 2017) and alter patterns of neurochemical release (Benner et al., 2018). Importantly, chronic doses of OT in rodents fail to produce anxiolytic effects normally seen with single doses (Du et al., 2017) which mirrors our present observations and suggests a translational mechanism of high clinical relevance. The apparent long-lasting desensitization effects of daily OT administration may be contributed to by the magnitude of the doses being administered and if so it is possible that lower daily doses might produce reduced effects.

The amygdala is one of the main neural substrates mediating OT’s functional effects and its attenuation of fear of reactivity in this region is considered as a core therapeutic mechanism of action (Jurek and Neumann, 2018, 2018; Kendrick et al., 2017). Importantly, we have additionally demonstrated that the magnitude of amygdala responses to fear faces are positively associated with intensity and arousal ratings in the PLC group and that OT exerts an anxiolytic action by reducing both these ratings and abolishing their correlation with amygdala activation. Dysregulations in amygdala and behavioral responses to fear stimuli have been observed across major psychiatric disorders, particularly those characterized by marked social impairments and anxiety (Hennessey et al., 2018; Neumann and Slattery, 2016). Furthermore, our finding that both neural and behavioral anxiolytic effects of OT are stronger after repeated compared to single doses in the group receiving OT every other day supports the assumption that optimal therapeutic effects should be obtained following chronic treatment.

Our second major finding is that OT’s attenuations of neural and behavioral responses to fear faces are highly dependent on OXTR genotype. Only G-non-carriers of rs53576 (i.e. AA) and A-carriers of rs2254298 (i.e. AA and AG) and AA homozygotes for rs53576 and A+ carriers of rs2254298 were significantly responsive to OT. A-carriers of both SNPs have frequently, although not universally, been associated with social dysfunction in autism as well as social anxiety (Cataldo et al., 2018; Jurek and Neumann, 2018). A recent haplotype-based analysis of OXTR SNPs including rs53576 and rs2254298 also indicated that individually they have some association with sensitivity to OT-effects on face recognition (Chen et al., 2015). While the current study clearly indicates that OT modulation of amygdala and visual cortex responses to fear are strongly associated with the AA genotype (Cohen’s *ds* from 0.81-1.41), it remains to be seen whether other OT-dependent effects exhibit the same association. Limitations to the current findings are that subject numbers are still relatively low for establishing robust genetic associations and that the AA allele of rs53576 occurs more frequently in Asian compared with Caucasian populations (Butovskaya et al., 2016). Also, rs53576 exhibits some sex-dependent differences in social cooperation effects of OT (Feng et al., 2015b) and our current study focused only on males to avoid potential menstrual cycle effect issues.

Interestingly, task-related and intrinsic network changes produced by OT showed a strikingly different sensitivity to repeated doses and OXTR genotype. However, importantly on the fifth day of treatment there was no advantage of giving OT daily as opposed to every other day. Intrinsic networks may be less influenced by repeated doses of OT or OXTR genotype than those activated by tasks, although arguably in the context of OT’s putative therapeutic effects its task-dependent impact of neural circuitry engaged during social interactions should be of greatest importance. Future studies should measure differential OT effects on both task-related and intrinsic networks.

In conclusion, the current study provides the first evidence for an important influence of dose frequency and receptor genotype on the neural and behavioral actions of intranasal OT in response to fear faces in healthy human subjects. Dose frequency therefore requires further empirical evaluation in patient populations given that it can critically determine treatment efficacy in clinical trials employing chronic administration as an intervention in psychiatric disorders.

## Material and Methods

### Study design

The main objectives of the study were to firstly investigate the effects of acute (single dose) and repeated doses (daily or every other day for 5 days) of 24IU OT versus PLC on two biomarkers: (1) amygdala and behavioral responses to fearful faces and (2) resting state functional connectivity between the amygdala and mPFC. Secondly we investigated modulatory influences of OXTR rs53576 and rs2254298 polymorphisms on sensitivity to intranasal OT.

#### Participants

A total of 147 healthy, right-handed healthy adult male subjects were enrolled according to common inclusion and exclusion criteria for human OT-administration studies. This subject number was determine a priori based on achieving 85% power for an expected medium effect size of 0.5. A total of 9 subjects were excluded due to failure to complete the study or excessive head movement (see Fig 1). Subjects were randomly assigned to repeated intranasal treatment (single daily dose on five consecutive days) of (1) placebo (PLC; n = 46, M ± SD, 22.46± 2.3 years), (2) oxytocin (OT_5;_ n = 49, M ± SD, 21.78 ± 2.3 years), or interleaved OT and PLC (OT on days 1, 3, 5, PLC on days 2, 4; OT_3_; n = 43, M ± SD, 21.02 ± 2.0 years) (see **SI**). To ensure compliance all subjects were required to come to the center every day and supervised during self-administration of nasal sprays. To control for between-group differences in potential confounders pre-treatment levels of anxiety, depression, and empathy and trait autism were assessed using validated scales. There were no significant differences between the OT and PLC groups (Table S4). The study was approved by the local ethics committee (Institutional Review Board, University of Electronic Science and Technology of China) and subjects provided written informed consent. The study was in accordance with the latest revision of the Declaration of Helsinki, pre-registered at Clinical Trials.gov (NCT03610919 - https://clinicaltrials.gov/ct2/show/NCT03610919) and in line with recommendations for trials in psychological experiments (Guidi et al., 2018) (see Fig. 1 for Consort flow diagram). Subjects received monetary compensation for participation.

### Experimental procedures

The study employed a double-blind, randomized, placebo-controlled, between-subject design. The OT and PLC sprays used in the 3 groups were both supplied by Sichuan Meike Pharmaceutical Co. Ltd, Sichuan, China in identical dispenser bottles containing identical ingredients (glycerine and sodium chloride) other than OT. In line with recommended guidelines experiments started 45 minutes after intranasal administration (Guastella et al., 2013). In post-treatment interviews collected on days 1 and 5 subjects were unable to identify better than chance whether they had received OT or PLC (χ^2^ < 0.1, ps > 0.2, Table S5) confirming successful blinding over the entire study period. For OXTR genotyping subjects provided buccal swaps on the 1^st^ day for analysis of OXTR rs2254298, rs53576 SNPs (see **SI** and (Montag et al., 2017)).

For the implicit face-emotion processing task 208 grayscale facial stimuli displaying happy, neutral, angry or fearful facial expressions (n = 26 per category, 50% female) were used. Stimuli were initially rated with respect to arousal and valence by an independent group of subjects and two matched independent sets of stimuli were produced for use on the 1^st^ and 5^th^ days (all ps > 0.3, see **SI** for details and Table S6). The presentation order of the two sets was counter balanced. To ensure attentive processing subjects were required to identify the gender of each face picture (see **Figure S2**).

### Primary outcomes and analysis plan

Face emotion-related amygdala responses were assessed using the event-related implicit face processing fMRI paradigm on treatment days 1 and 5. Intrinsic amygdala connectivity was assessed by means of two resting state fMRI assessments (before and after the task-paradigm). Valence, arousal and intensity ratings (scale: 1-9) for the facial stimuli were collected immediately after MRI acquisition as additional behavioral outcomes (further details see **SI**). In line with the main aim of the study changes in amygdala fear reactivity and amygdala intrinsic connectivity between the 1^st^ and 5^th^ day served as primary outcome measures. Changes in valence, arousal and intensity ratings for the fear faces represented an associated behavioral outcome measure.

### Imaging acquisition and analysis

MRI data was acquired using standard sequences on a 3T GE MR750 system. In addition to the functional time series high resolution T1-weighted structural MRI data was acquired at both time points to improve normalization and control for acute and chronic effects of treatment on brain structure. MRI data was preprocessed using validated procedures in SPM12 (Friston et al., 1994) (Statistical Parametric Mapping; http://www.fil.ion.ucl.ac.uk/spm) and Data Processing Assistant or Resting-State fMRI(Yan, 2010) (DPARSFA; http://rfmri.org/DPARSF) (for details see **SI**). First level General Linear Models (GLM) for the task-related fMRI data included separate regressors for the four emotional conditions, gender identity rating period and 6 movement parameters and appropriate contrasts were subjected to a second level random effects analysis.

### Statistical analyses

To identify OT-sensitive regions and confirm previous findings on suppression of amygdala reactivity following single-dose OT-administration (Young and Barrett, 2015) a voxel-wise whole-brain two-sample t-test was conducted in SPM comparing neural activity in OT- and PLC-treated subjects on the1^st^ day. For all subsequent analyses a 6-mm sphere centered at the maximum t-values of the acute OT effects in the amygdala served as target region to determine different trajectories of amygdala functioning following acute versus repeated treatment (for similar approach see (Spengler et al., 2017)).

### Primary outcomes: amygdala and behavioral fear-face reactivity and intrinsic connectivity

Differences in stimulus-induced amygdala and behavioral responses to fear faces were examined by means of mixed ANOVAs with treatment (PLC, OT_3,_ OT_5_) as between-subject factor, time point (1^st^ day, acute effects; 5^th^ day, chronic effects) as within-subject factor. Dependent variables were extracted target region amygdala-reactivity towards (parameter estimates extracted using Marsbar, http://marsbar.sourceforge.net) and subsequent valence, intensity and arousal ratings of fearful faces. Bonferroni-corrected post-hoc comparisons were used to explore significant interactions. Associations between amygdala activation in response to fear faces and subsequent behavioral ratings of their valence, intensity and arousal were assessed using Pearson correlation and group differences calculated using Fishers-z tests.

Differences in resting state amygdala functional connectivity were determined using a voxel-wise seed-to-whole-brain ANOVA as implemented in the SPM flexible factorial design with treatment (PLC, OT_3,_ OT_5_) as between-subject factor, time point (1^st^ day, acute effects; 5^th^ day, chronic effects) as within-subject factor and amygdala connectivity maps as dependent variable. To account for potential effects of task-engagement on intrinsic amygdala connectivity the primary analysis focused on pre-task resting state maps. A subsequent ROI analysis on the post-task data (focusing on extracted estimates from a 6mm sphere centered at vmPFC determined by the pre-task data) served as a replication and further validation.

### Secondary outcome measures: influence of OXTR genotype

For rs2254298 subjects were divided into A+ carriers (AA and AG) and A-non-carriers (GG) and rs53576 into G+ carriers (GG and GA) and G-non-carriers (AA). To explore effects of genotype as a secondary outcome measure, OXTR group was included as additional between-subject factor in the corresponding ANOVAs and amygdala fear reactivity, amygdala-vmPFC intrinsic functional connectivity and behavior rating response served as dependent variables.

### Control for treatment effects on brain structure

To control for potential confounding effects of single- and repeated OT-administration on brain structure, a voxel-based morphometry (VBM) analysis was conducted on the T1-weighted images acquired on both testing days using SPM12 standard procedures (Ashburner, 2007; Ashburner and Friston, 2005). Effects of single- and repeated dose administration were explored concordant with the fMRI analyses (see **SI**).

### Thresholding and statistical implementation

Voxel-wise whole brain analyses in SPM were thresholded at a cluster-based Family-wise error (FWE) corrected level (p < .05) with an initial cluster forming threshold of p < .001 (in line with recent recommendations (Eklund et al., 2016; Mueller et al., 2017)). Behavioral and neural (extracted parameter estimates) indices were analyzed using SPSS 22.0 and appropriate ANOVA models. Partial eta squared (*η*^2^_p_) and Cohen’s *d* were computed as measures of effect size. Group differences in parameter estimates extracted from significant main effect in SPM were further evaluated using two-sample t tests. All reported p values were Bonferroni-corrected, two-tailed, and p ≤ 0.05 considered significant.

## Acknowledgement

We thank the reviewers and editor for their comments and suggestions for improving the manuscript.

## Author contributions

JK and KMK designed the experiment. JK, YZ and CS carried out the experiment. JK, KMK, CM, FZ and BB analyzed the experiment and JK, KMK, CM and BB wrote the paper. All authors contributed to the conception of the study and approved the paper.

## Supplementary information for

### Supplementary Methods

#### Participants

147 healthy, right-handed healthy adult subjects were recruited by local advertisement. All subjects had normal or corrected to normal vision and no history of or current neurological or psychiatric disorders. Participants were free from current (one month before the experiment) or regular use of medication and were required to abstain from caffeine, alcohol and tobacco at least 24 hours before the experiment. Four subjects were not able to attend all study appointments and 5 were excluded due to head movement during the fMRI assessment (motion > 3.0mm translation or 3°and mean frame-wise displacement (FD < 0.5 mm) (Power et al., 2012), Pre-treatment levels of anxiety, depression, trait autism and empathy were assessed using validated scales (State-Trait Anxiety Inventory (STAI) (Kvaal et al., 2005), Beck Depression Inventory (Beck et al., 1988), Autism-Spectrum Quotient (AQ) (Baron-Cohen et al., n.d.), and Interpersonal Reactivity Index (IRI) (Siu and Shek, 2005).

#### Stimuli and face task

208 gray scaled facial stimuli displaying happy, neutral, angry or fearful facial expressions (n = 26 per category, 50% female) were used employed for the implicit emotional face processing task. Stimuli were initially rated with respect to arousal and valence by an independent group of participants (n = 24 participants, 10 females, age, M ± SD, 21.3 ± 1.9 years). Based on these valence and arousal ratings two independent sets of stimuli for the 1^st^ day and 5^th^ day were constructed with matched arousal and valence (all ps > 0.3, see Table S6) and the order of the two sets are counter balanced in the treatment group and control group.

Subjects were instructed to attentively process the presented facial stimuli and were required to indicate the gender of the actor after presentation (left button press for male, right button press for female face) to ensure attentive processing. Stimuli were presented for 3s followed by a jittered fixation (1.5-2.1s) followed by the gender discrimination task presented for 2s (2s) and followed by a jittered inter-trial interval (4.5-6s) that served as low level baseline. Order of gender and emotion was pseudo-randomized (see also Fig. S2).

#### Imaging acquisition of fMRI data

fMRI data employing blood oxygenation level-dependent (BOLD) contrast was acquired on a 3T GE MR750 system. Functional images for task-based and resting state data assessments were acquired using EPI sequences (TR=2000ms, echo time=30ms, flip angle=90°, FOV=240mm × 240mm, voxel size= 3. 75 × 3. 75 ×4 mm, resolution=64 × 64, the number of slices=39). T1 -weighted anatomical images were additionally acquired to improve normalization of the functional images and additionally served to explore structural changes related to OT treatment (TR=6ms, echo time=2ms, flip angle=9°, FOV=256mm × 256mm, voxel size =1×1×1mm, number of slices=156).

#### fMRI data preprocessing (Implicit emotional face processing task part)

Task-related fMRI data analyzed using SPM12 (Friston et al., 1994) (Statistical Parametric Mapping; http://www.fil.ion.ucl.ac.uk/spm). The first 5 volumes were discarded to allows T1 equilibration and the remaining images were applied to correct for temporal slice acquisition differences and then realigned and unwarped to correct for head motion (six-parameter rigid body algorithm), and slice-timing correction.

Tissue segmentation, bias-correction were applied to the high-resolution structural images. After co-registration of the functional time-series with the skull-stripped anatomical scan the transformation matrix was applied to normalized the functional images to MNI space with a voxel size of 3 × 3 ×3 mm. Normalized images were spatially smoothed using a Gaussian kernel with full-width at half-maximum (FWHM) of 5 mm. Participants with head motion exceeding 3.0mm translation or 3°rotation and mean frame-wise displacement (FD > 0.5 mm) were excluded. Mean frame-wise displacement (FD) were controlled between OT treatment group and PLC control group (see Table S7)

#### fMRI data preprocessing (resting state)

Resting state data was analyzed using SPM12 (Statistical Parametric Mapping; http://www.fil.ion.ucl.ac.uk/spm) and Data Processing Assistant for Resting-State fMRI(Yan, 2010) (DPARSFA; http://rfmri.org/DPARSF). After discarding the first 5 scans, preprocessing included slice-time correction, head motion correction using six-parameter rigid body algorithm. Then they were co-registered with the anatomical scan and normalized to MNI space with voxel size of 3 × 3 ×3 mm. Normalized images were then spatially smoothed using a Gaussian kernel with full-width at half-maximum (FWHM) of 5mm. As for nuisance signal correction, the following nuisance parameters were included as regressors, 6 head motion parameters, 6 head motion parameters one time point before, and their squares, white matter (WM) and cerebrospinal fluid (CSF). After filtering with the band of (0.01-0.1HZ) and linear detrending, the functional connectivity of data has been calculated on the whole brain level based on the region of activation during task. Participants with head motion exceeding 3.0mm translation or 3°rotation and mean frame -wise displacement (FD > 0.5 mm) were excluded. Mean frame-wise displacement (FD) were controlled between OT treatment and PLC control groups (see Table S7).

#### fMRI data preprocessing (voxel-based morphometry part)

To further control for potential confounding effects of single- and repeated dose administration of OT on brain structure a voxel-based morphometry (VBM) analysis was performed on the T1-weighted images acquired on both testing days. Processing for longitudinal effects included unified segmentation, DARTEL preprocessing, and Pairwise Longitudinal Registration (Ashburner, 2013) using in SPM12 according to standard procedures(Ashburner, 2007; Ashburner and Friston, 2005). Repeated administration effects on gray matter (GM) were assessed using whole-brain analysis across groups. For the acute effect identical preprocessing was applied without the longitudinal registration step including T1 images from the 1^st^ day and an independent samples t test to compare OT and PLC treated participants.

#### Genotyping

A total of 120 subjects (samples could not be analyzed from 27 subjects either due to problems with buccal cell sampling or equivocal genotyping results) were successfully genotyped from buccal cell swabs. DNA was extracted by a MagNa Pure 96 robot (Roche Diagnostics, Mannheim, Germany) with a commercial Viral NA Small Volume Kit (Roche Diagnostics, Mannheim, Germany). Polymorphisms were analyzed by means of a real time quantitative polymerase chain reaction (PCR) and subsequent high resolution melting detection analysis with a LightCycler Cobas z480 (Roche Diagnostics, Mannheim, Germany). Simple probe assay designs provided by TibMolBiol (Berlin, Germany) were conducted. The following SNPs were investigated for participants in the present study: OXTR rs2254298, rs53576 (see Tables S2 **and** S3 for distribution and Hardy Weinberg details).

### Supplementary Results

#### Bilateral calcarine cortex responses to fear faces and associations with OXTR genotype

Mixed ANOVAs with treatment (PLC, OT_3,_ OT_5_) and time point (1^st^ day, acute effects; 5^th^ day, chronic effects) as factors revealed a significant main effect of treatment (F_2, 135_ =11.52, p < 0.001, *η*^2^_p_ =0.146) and treatment x time point interaction (F_2, 135_ = 3.22, p=.043, *η*^2^_p_ = 0.046). Post hoc comparisons confirmed that OT significantly decreased calcarine cortex responses in the OT_3_ (p < 0.001, *d* = 0.97) and OT_5_ (p < 0.001, *d* = 0.72) groups relative to PLC on the1^st^ day. No differences were observed between the OT-treatment groups on the 1^st^ day (p = 0.351, *d* = 0.21). On the 5^th^ day, suppression of calcarine cortex fear reactivity relative to the PLC-group was maintained in the OT_3_ (p < 0.001, *d* = 0.81) but not the OT_5_ group (p = 0.326, d = 0.19) and the OT_3_ group was also significantly different from the OT_5_ group (p = 0.006, *d* = 0.62). Thus, the bilateral calcarine cortex showed similar evidence to the right amygdala for a reduced effect of OT with daily repeated doses after 5 days.

For calcarine cortex responses there was a significant treatment x genotype interaction for rs53576 (F_2, 105_= 5.17, p = 0.007, *η*^2^_p_ = 0.09) although for rs2254298 it did not reach significance (F_2, 105_= 2.18, p = 0.118, *η*^2^_p_ = 0.04). For rs53576, OT-induced suppression on day 1 was only significant in G-non-carriers (AA – p <0.001, Cohen’s *d* =1.12) and also on the 5^th^ day in the OT_3_ group (AA - p < 0.001, Cohen’s *d* =1.28) and OT_5_ group (p = 0.006, Cohen’s *d* =0.92). An exploratory post hoc analysis for rs2254298 revealed that OT-induced suppression on the 1^st^ day across treatment groups was only significant in A-carriers (AA and AG-p < 0.001, Cohen’s *d* =1.20) and also on the 5^th^ day in the OT_3_ group (AA and AG - p < 0.001, Cohen’s *d* =1.24). Thus the calcarine cortex showed a similar pattern of receptor genotype dependency for the effects of OT as the amygdala.

#### Bilateral calcarine cortex responses to fear faces and associations with intensity and arousal ratings

Associations between calcarine (visual) cortex responses to fear faces and arousal and intensity ratings were marginal in the PLC group on the 1^st^ day (arousal: r = 0.28, p = 0.059; intensity: r = 0.22, p = 0.150) and were not influenced by OT (arousal: OT_3_, r = 0.15 p = 0.329; Fisher’s z = 0.31, p = 0.540; OT_5_, r = 0.08, p = 0.584; z = 0.98, p = 0.328; intensity: OT_3_, r = 0.24, p = 0.123; Fisher’s z = 0.11, p = 0.912; OT_5_, r = 0.12, p = 0.423; z = 0.18, p = 0.859). However, on the 5^th^ day there was a significant association for arousal ratings in the PLC group (PLC: r = 0.35 p = 0.016) and this was significantly reduced in the OT_3_ group (r = −0.10 p = 0.544; z = 2.12, p = 0.034) but not the OT_5_ group (r = 0.05 p = 0.725; z = 1.50, p = 0.134). Intensity ratings on the 5^th^ day showed a similar pattern (PLC: r = 0.37 p = 0.012; OT_3_, r = −0.05 p = 0.734; z = 1.98, p = 0.047; OT_5_ group, r = 0.108 p = 0.462; z = 1.29, p = 0.195) (Fig.S4).

#### Superior frontal gyrus responses to fear faces and associations with OXTR genotype

Mixed ANOVAs with treatment (PLC, OT_3,_ OT_5_) and time point (1^st^ day, acute effects; 5^th^ day, chronic effects) as factors revealed a significant main effect of treatment (F_2, 135_ = 6.80, p = 0.002, *η*^2^_p_ =0.092) but no treatment x time point interaction (F_2, 135_=0.55, p = 0.576, *η*^2^_p_ =0.008). Exploratory post hoc comparisons confirmed that OT significantly decreased superior frontal gyrus responses in the OT_3_ (p = 0.005, *d* = 0.59) and OT_5_ (p = 0.001, *d* = 0.67) groups relative to PLC on the 1^st^ day. No differences between the OT-treatment groups were observed on the 1^st^ day (p = 0.753, *d* = 0.06). On the 5^th^ day, suppression of superior frontal gyrus fear reactivity relative to the PLC-group was maintained in the OT_3_ group (p = 0.011, *d* = 0.63) and the OT_5_ group (p = 0.045, *d* = 0.38). No differences between the OT-treatment groups were observed on the 5^th^ day (p = 0.528, *d* = 0.13). Thus, there was no evidence for reduced superior frontal gyrus responses to daily OT doses being reduced after 5 days.

For superior frontal gyrus responses there was no significant treatment x genotype interaction for rs53576 (F_2, 105_= 1.19, p = 0.309, *η*^2^_p_ = 0.02) or for rs2254298 (F_2, 105_= 1.28, p = 0.279, *η*^2^_p_ = 0.02) indicating that there was no significant influence of OXTR genotype on responses.

#### Superior frontal gyrus responses to fear faces and associations with arousal and intensity ratings

There were no significant associations between superior frontal gyrus responses to fear faces and arousal and intensity ratings for PLC or OT groups on the 1^st^ or 5^th^ days (ps > 0.11) (Fig.S5).

#### Neural and behavioral responses to fearful faces in association with combined rs53576 and rs2254298 genotypes

We also carried out an exploratory analysis of neural responses and intensity ratings for fearful faces for the 4 different combined rs53576 and rs2254298 genotypes following acute OT treatment (i.e. using combined OT_3_ and OT_5_ groups on the 1^st^ day versus PLC to increase power). The 4 different combined genotypes had an approximately equal distribution (see Fig. S3). For right amygdala responses, there was a marginal significant treatment x genotype interaction for combined genotype of rs53576 and rs2254298 (F_2, 110_= 2.20, p = 0.093, *η*^2^_p_ = 0.06). Post hoc analysis indicated OT-induced suppression on day 1 was significant in the group with both G-non-carriers of rs53576 (i.e. AA) and A-carriers of rs2254298 (i.e. AA or AG - p < 0.001, Cohen’s *d* =1.25) (see Fig. S3).

For the bilateral calcarine cortex response to fearful faces, the treatment x genotype interaction for the combined genotype of rs53576 and rs2254298 was marginally significant (F_2, 110_= 2.10, p = 0.105, *η*^2^_p_ = 0.06). Again an exploratory post hoc analysis indicated OT-induced suppression on the 1^st^ day was significant in the group with both G-non-carriers of rs53576 (AA) and A-carriers of rs2254298 (AA or AG - p = 0.001, Cohen’s *d* =1.26), similar to the pattern in right amygdala (see Fig. S3).

For the superior frontal gyrus responses there was no treatment x genotype interaction for the combined genotype of rs53576 and rs2254298 (F_2, 110_= 0.09, p = 0.96, *η*^2^_p_ = 0.003) and no significant exploratory post-hoc findings.

For intensity ratings of fearful faces there was a marginal treatment x genotype interaction for the combined genotype of rs53576 and rs2254298 (F_2, 110_= 2.35, p = 0.08, *η*^2^_p_ = 0.009) and an exploratory post-hoc analysis showed that OT-induced reductions in intensity ratings on the 1^st^ day were significant in the group with both G-non-carriers of rs53576 (AA) and A-carriers of rs2254298 (AA or AG - p = 0.009, Cohen’s *d* = 0.85). For arousal ratings for fearful faces there was no significant treatment x genotype interaction for the combined genotype of rs53576 and rs2254298 (F_2, 110_= 1.92, p = 0.13, *η*^2^_p_ = 0.053) but an exploratory post-hoc analysis showed that OT-induced reduction in intensity ratings on the 1^st^ day was marginally significant (after correction) in the group with both G-non-carriers of rs53576 (AA) and A-carriers of rs2254298 (AA or AG - p = 0.020, Cohen’s d=0.75) (see Fig. S3).

---

**Table S1.** Whole brain acute effect (1st day) of intranasal OT on neural responses to fearful faces

| Regions | x | y | z | k | P<sub>FWE</sub> |
| --- | --- | --- | --- | --- | --- |
| **OT<PLC (fear faces)** |  |  |  |  |  |
| Calcarine_R (aal) | 18 | -82 | 8 | 156 | 0.002 |
| Calcarine_L (aal) | 0 | -88 | 5 |  |  |
| Occipital_Sup_R (aal) | 18 | -91 | 20 |  |  |
| Frontal_Mid_R (aal) | 27 | 8 | 56 | 82 | 0.036 |
| Frontal_Sup_R (aal) | 33 | -4 | 59 |  |  |
| Frontal_Sup_R (aal) | 21 | -1 | 52 |  |  |
| Amygdala_R (aal) | 27 | -4 | -13 | 78 | 0.043 |
| Putamen_R (aal) | 30 | 5 | -7 |  |  |

**Table S2.** The number of A carriers (A+) and A non-carriers (A-) of rs2254298, G carriers (G+) and G non-carriers (G-) of rs53576 in each treatment group

| Genotypes | PLC | OT<sub>3</sub> | OT<sub>5</sub> | In total | $\chi^2$ | P | $\eta^2$ |
| --- | --- | --- | --- | --- | --- | --- | --- |
| rs2254298(A+) | 20 | 23 | 20 | 63 | 0.088 | 0.957 | 0.025 |
| rs2254298(A-) | 16 | 18 | 14 | 48 |  |  |  |
| rs53576(G+) | 15 | 17 | 20 | 52 | 2.824 | 0.244 | 0.124 |
| rs53576(G-) | 21 | 24 | 14 | 59 |  |  |  |

**Table S3.** Distribution of Genotypes in the Sample of N=120 participants

|  | Genotypes |  |  |  | HWE |
| --- | --- | --- | --- | --- | --- |
|  | AA: | AG: | GG: | Total: |  |
| rs2254298 | 15 | 53 | 52 | 120 | Chi<sup>2</sup> = 0.07,<br>*p* = .794 |
| rs53576 | 60 | 45 | 15 | 120 | Chi<sup>2</sup> = 1.94,<br>*p* = .163 |

**Table S4.** Pre-treatment anxiety, depression, autism and empathy scores in both groups

| Questionnaire | PLC | OT<sub>3</sub> | OT<sub>5</sub> | F | P |
| --- | --- | --- | --- | --- | --- |
| STAI, state | 38.78±11.53 | 35.49±10.69 | 37.14±8.92 | 1.13 | 0.33 |
| STAI, trait | 41.67±11.60 | 40.28±10.51 | 39.20±8.94 | 0.81 | 0.15 |
| BDI | 9.84±9.13 | 8.93±8.08 | 8.00±7.45 | 0.57 | 0.57 |
| AQ | 20.91±5.57 | 20.35±5.91 | 19.47±4.57 | 0.88 | 0.42 |
| IRI | 50.59±9.84 | 53.02±8.74 | 49.39±9.81 | 1.72 | 0.18 |

**Table S5.** Chi Squared Test of post-experiment interviews where subjects were required to identify which treatment they received. In all cases subjects were unable to guess better than chance.

| Day | PLC | OT<sub>3</sub> | OT<sub>5</sub> | In total | $\chi^2$ | P | $\eta^2$ |
| --- | --- | --- | --- | --- | --- | --- | --- |
| 1 | 24 | 20 | 22 | 66 | 0.547 | 0.76 | 0.062 |
| 2 | 20 | 15 | 17 | 52 | 0.988 | 0.61 | 0.078 |
| 3 | 24 | 28 | 24 | 76 | 2.645 | 0.27 | 0.014 |
| 4 | 19 | 18 | 20 | 57 | 0.01 | 0.99 | 0.003 |
| 5 | 25 | 25 | 24 | 74 | 0.787 | 0.68 | 0.039 |

**Table S6.** Ratings for each of the two sets of different face emotion stimuli used on the 1^st^ day and 5^th^ day in a counterbalanced design. There were no significant differences between the two sets for valence and arousal ratings.

|  | Valence |  |  |  | Arousal |  |  |  |
| --- | --- | --- | --- | --- | --- | --- | --- | --- |
|  | Set A | Set B | t | p | Set A | Set B | t | p |
|  | On the 1<sup>st</sup> day | On the 5<sup>th</sup> day |  |  | On the 1<sup>st</sup> day | On the 5<sup>th</sup> day |  |  |
| Happy | 6.26±0.94 | 6.25±0.97 | 0.04 | 0.97 | 5.50±0.59 | 5.52±0.44 | 0.12 | 0.90 |
| Neutral | 5.67±0.17 | 5.61±0.16 | 1.06 | 0.30 | 3.03±0.36 | 2.98±0.32 | 0.55 | 0.59 |
| Fear | 6.40±1.02 | 6.31±0.17 | 0.34 | 0.74 | 6.71±0.70 | 6.70±0.63 | 0.03 | 0.98 |
| Angry | 6.16±0.98 | 6.04±1.04 | 0.48 | 0.64 | 5.66±0.84 | 5.83±0.99 | 0.82 | 0.42 |

**Table S7.**
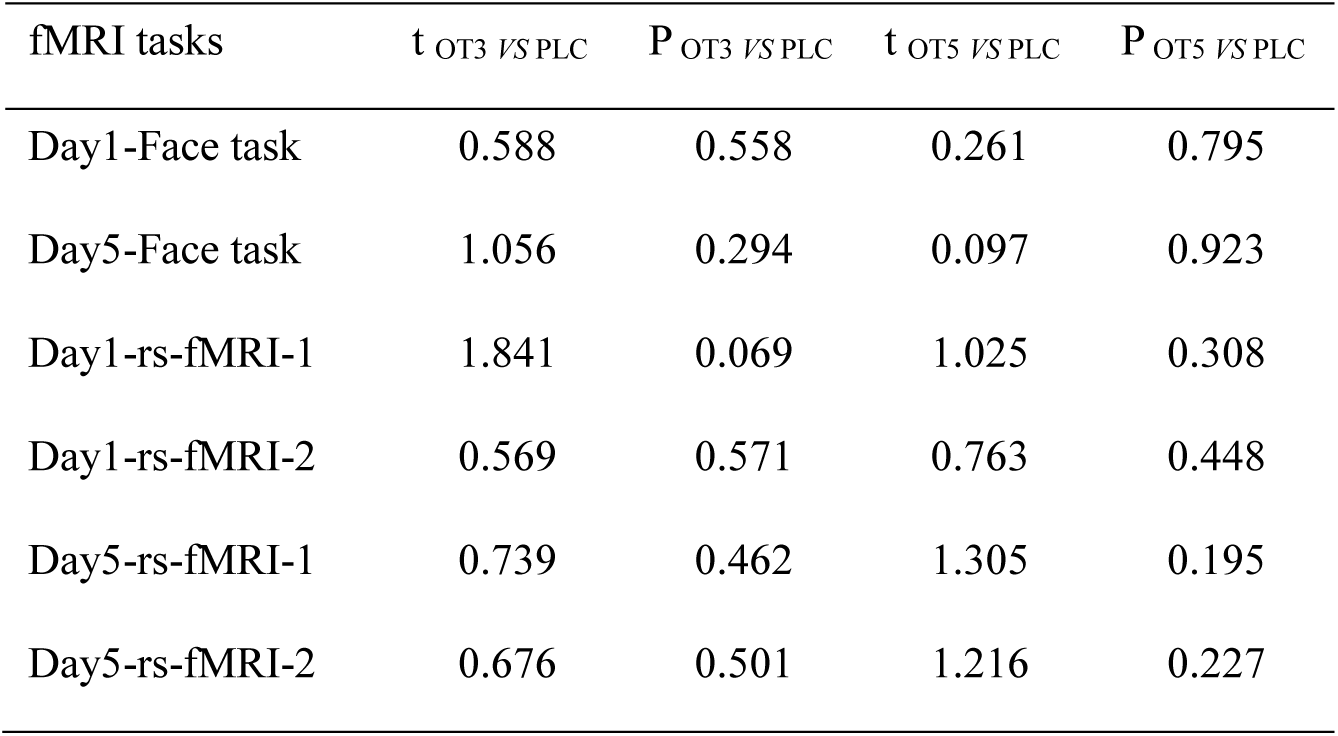
Group Comparison of Mean frame-wise displacement

**Fig.S1.**
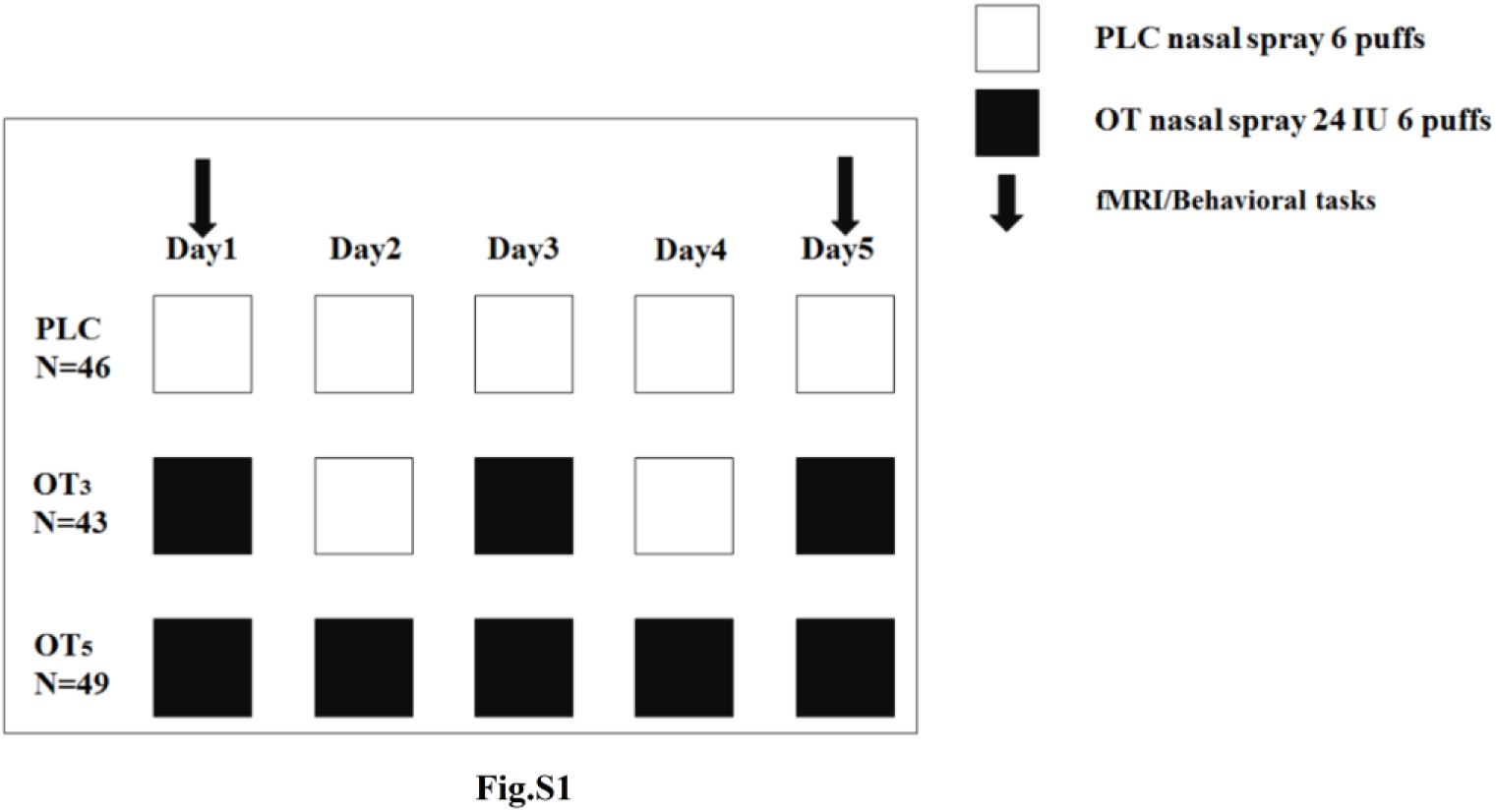
Treatment protocol. Participants were randomly assigned to have nasal spray of oxytocin (OT) for 5 days 24 IU per day (OT_5_ group) or have OT or placebo nasal spray on alternate days during the 5 days (OT on the 1^st^, 3^rd^ and 5^th^ day), 24 IU per day (OT_3_ group) or have daily nasal spray of PLC for 5 days (PLC group).

**Fig.S2.**
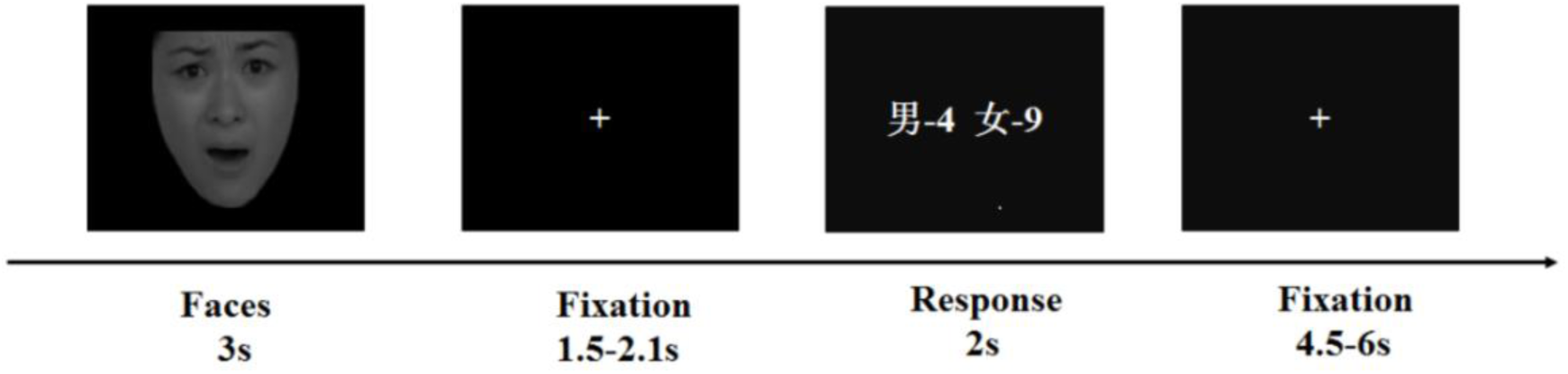
Face emotion task procedure. Face stimuli appeared on the screen for 3 seconds, and then on the subsequent response screen participants were required to press the left or right response key to identify the gender of the face they had seen.

**Fig.S3.**
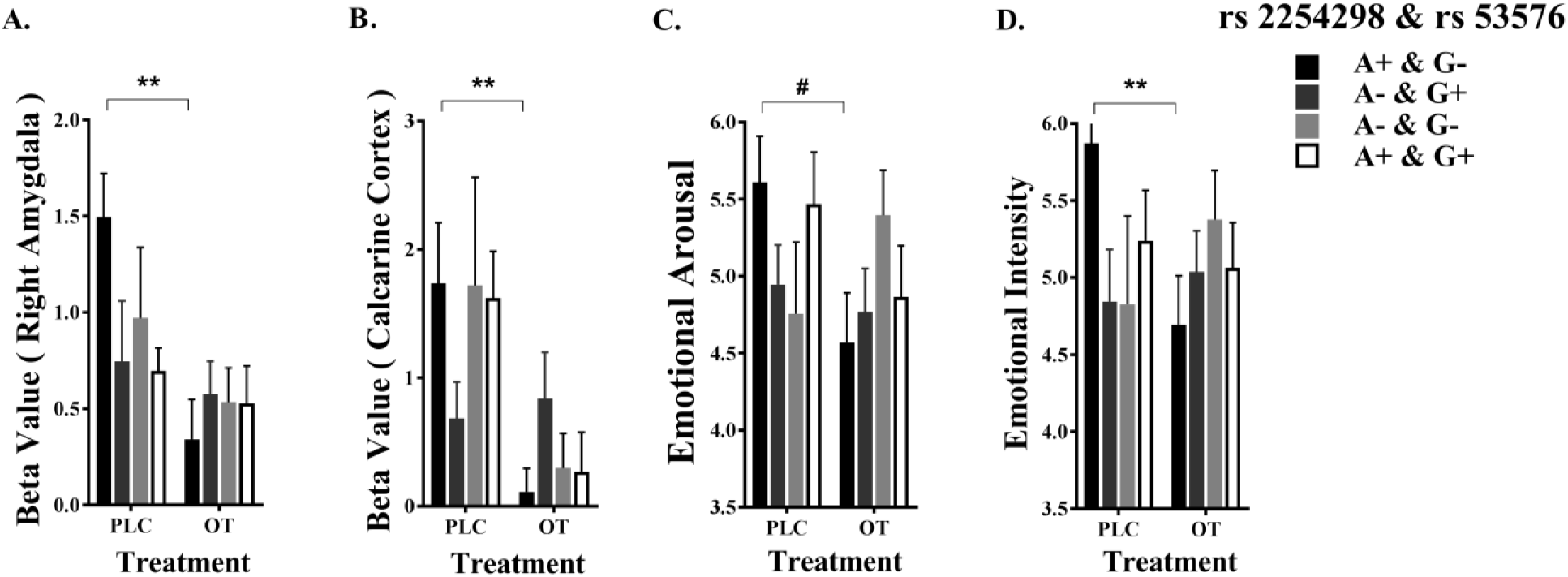
The influence of combined rs53576 and rs2254298 genotypes on oxytocin acute treatment effect on brain activation and behavioral response (A) right amygdala and (B) bilateral calcarine gyrus responses to fear faces in the placebo (PLC) and combined oxytocin (OT) treatment groups on the 1^st^ treatment day and (C) the same for arousal ratings (D) and intensity ratings of fear faces. ** p <0.001 and # p <0.05. After Bonferroni correction p <0.0125 considered significant. Subject numbers for combined rs 2254298 and rs 53576: A+ & G-13 (PLC), 24 (OT); A- & G+ 8 (PLC), 18 (OT); A- & G-8 (PLC), 14 (OT); A+ & G+ 7 (PLC), 19 (OT).

**Fig.S4.**
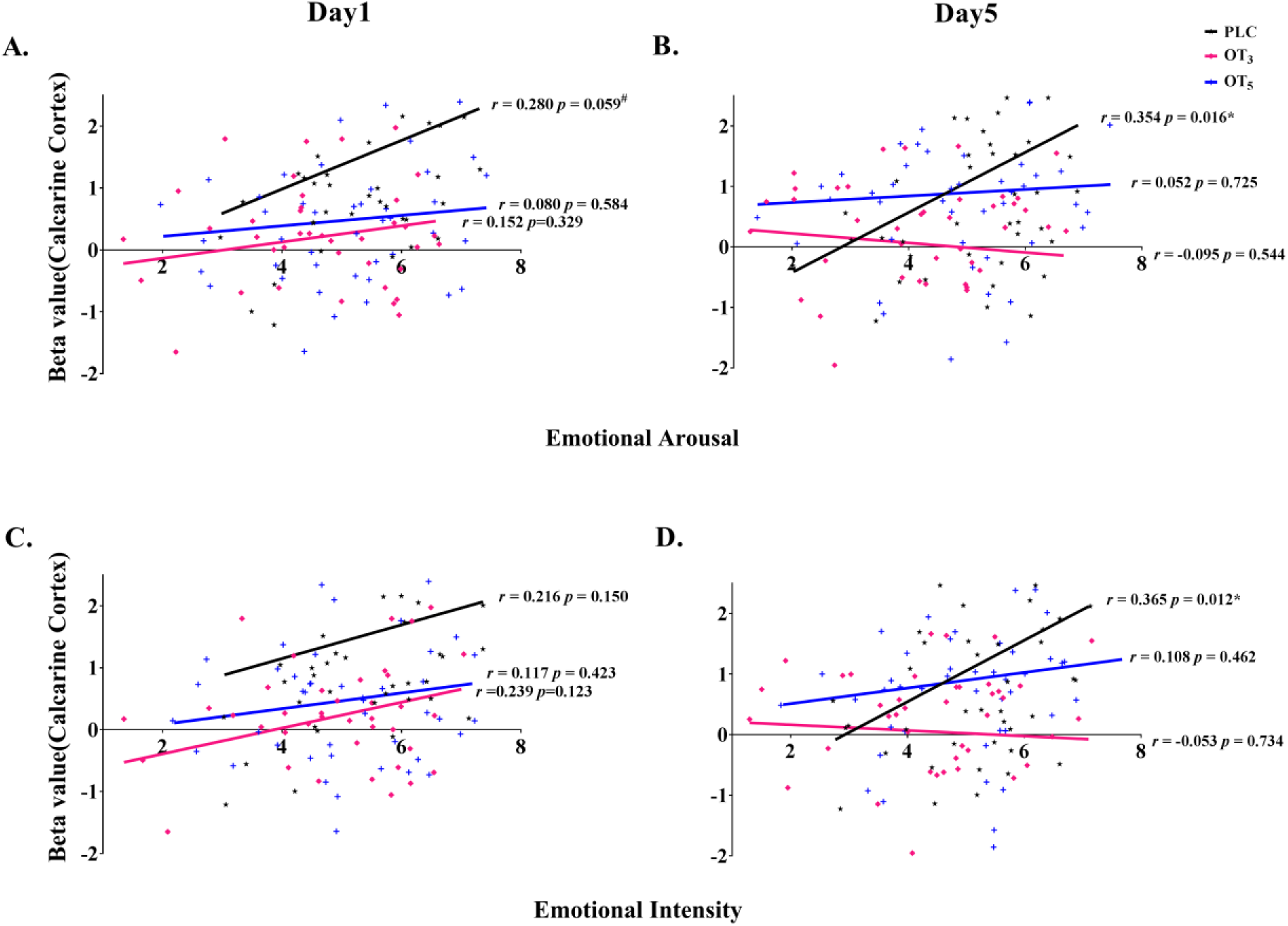
Associations between calcarine cortex (visual cortex) responses to fear faces and arousal and intensity ratings in the three treatment groups (PLC, OT_3_ and OT_5_). * p <0.05. # p<0.1.

**Fig.S5.**
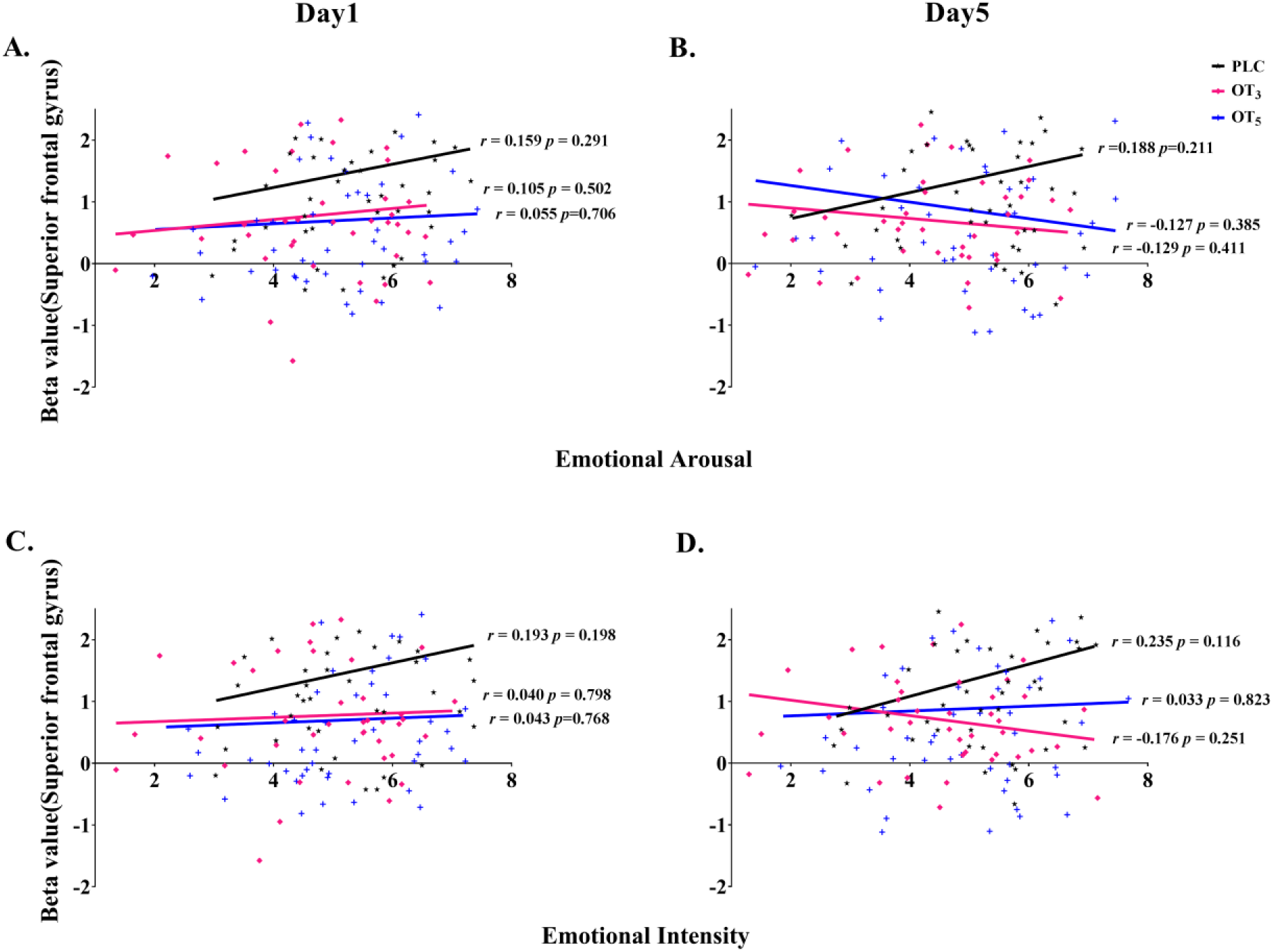
Associations between superior frontal gyrus responses to fear faces and arousal and intensity ratings in the three treatment groups (PLC, OT_3_ and OT_5_). * p <0.05

